# Predicting Disease-gene Associations through Self-supervised Mutual Infomax Graph Convolution Network

**DOI:** 10.1101/2023.07.13.548865

**Authors:** Jiancong Xie, Jiahua Rao, Junjie Xie, Yuedong Yang

**Affiliations:** School of Computer Science and Engineering at Sun Yat-sen University; School of Data and Computer Science at the Sun Yat-Sen University

**Keywords:** Mutual information, graph convolution network, disease-gene associations prediction

## Abstract

Illuminating associations between diseases and genes can help reveal the pathogenesis of syndromes and contributes to treatments, but a large number of associations still remained unexplored. To identify novel disease-gene association, many computational methods have been developed using disease and gene related prior knowledge. However, these methods remain of relatively inferior performance due to the limited external data sources and the inevitable noise among the prior knowledge. In this study, we have developed a new method, Self-Supervised Mutual Infomax Graph Convolution Network (MiGCN), to predict disease-gene associations under the guidance of external disease-disease and gene-gene collaborative graphs. The noises within the collaborative graphs were eliminated by maximizing the mutual information between nodes and neighbors through a graphical mutual infomax layer. In parallel, the node interactions were strengthened by a novel informative message passing layer to improve the learning ability of graph neural network. The extensive experiments showed that our model achieves performance improvement over the state-of-art method by more than 8% on AUC. The datasets, source codes and trained models of MiGCN are available at https://github.com/biomed-AI/MiGCN.

## 1 Introduction

Identifying the complex associations between diseases and genes is important for understanding the underlying mechanisms of diseases and contributing to the development of therapeutic treatments [1, 2]. Thus, the accurate identification of these associations is becoming one of the major biomedical research priorities. In the past decade, much effort has been devoted to building disease-gene associations databases, such as the Online Mendelian Inheritance in Man (OMIM) [3]. However, these databases suffer from the high sparsity of disease-gene data due to the costly and time-consuming information obtained through wet experiments.

In order to identify the unknown disease-gene associations, many computational methods have been developed. In early year, semi-supervised learning (SSL) methods were widely used [4, 5], such as label propagation approaches and positive-unlabeled learning. These SSL based classifiers were learned from the labeled (known disease genes) and unlabeled (the remaining genes) sets. Since the performance of these methods were limited by the naive model and single biological data source, deep learning methods were applied by incorporating disease and gene related information, such as disease phenotype similarity [6], gene expression matrix [7] and gene function interaction network [8]. For instance, IMC[9] constructed gene and disease feature vectors using gene-expression levels and disease text-mining information and integrated the features into a matrix factorization framework. The prior knowledge of diseases and genes can provide diverse information and a multi-view perspective and help deep learning models to better predict disease-gene associations. However, such feature-based approaches rely heavily on feature engineering with limited generalizability.

Since biological molecules perform their functions in corresponding pathways in a collaborative fashion, projecting their specific roles and collaborations onto networks or graphical structures can reveal more useful knowledge and provide more systematical aspects. Recently graph-based methods prevail, where a heterogeneous graph is usually constructed with diseases and genes as nodes and the associations as edges. For examples, CATAPULT[10] constructed a comprehensive network by integrating phenotype-gene network from eight other species and computed the closeness between candidate genes and known disease genes through random walking. GCN-MF[11] further computed gene-gene and disease-disease similarity graphs and employed graph convolution network (GCN[12]) to learn the non-linear representations of diseases and genes.

Albeit powerful, these methods still have some limitations. Firstly, previous graph-based models did not have enough communications between nodes. Indeed, the performance of GCN is strongly dependent on sufficient message passing as indicated in recent studies[13, 14]. Secondly, these methods haven’t fully used the prior knowledge of diseases and genes. In fact, with the accumulation of more and more gene sequencing and annotation, as well as disease phenotype data, many related databases have recently been well culled, such as PharmKG[15], GNBR[16]. These novel databases can provide more comprehensive and reliable information for disease-gene associations prediction. In addition, previous methods are hurt by the inevitable noises due to their direct use of the knowledge database.

Recently, self-supervised graph structure learning has been shown able to preserve various sources of knowledge while able to eliminate the noises. Many studies focus on the idea of mutual information estimation to learn an optimal node representation in an unsupervised manner[17, 18]. Intuitively, the mutual information of two nodes connected in the graph is greater than that of unconnected nodes. Maximizing the mutual information between a node and its neighbors enables the node to contain more information of its neighbors and to eliminate redundant information.

In this study, inspired from the self-supervised learning strategy, in this study, we have developed a new method, the Self-Supervised Mutual Infomax Graph Convolution Network (MiGCN), to identify disease-gene associations with the guidance of the gene-gene and disease-disease collaborative graphs. Specifically, we construct two collaborative graphs from external gene-gene interactions and disease-disease associations information, which are individually input into a self-supervised mutual infomax module to learn the node embeddings by maximizing mutual information between nodes and neighbors through negative sampling strategy. The learned embeddings and the graphs are further input into an informative message passing learning module to exploit the graph information for identifying new disease-gene associations. By evaluated on the OMIM dataset, the proposed model was shown to outperform state-of-the-art approaches with a large margin.

## 2 Materials and methods

The problem of prediction disease-gene associations is treated as a graph link prediction task. Each node represents a disease or gene, and each edge represents the disease-gene association. Our goal is to predict the potential links between disease nodes and gene nodes.

### 2.1 Datasets

#### 2.1.1 Disease-gene Associations data

We obtained human disease-gene association data from The Online Mendelian Inheritance in Man (OMIM) database[3], which contained 3954 observed phenotypes-gene associations spanning 3209 disease phenotypes and 12231 genes. The disease-gene association matrix is extremely sparse, with >90% of the columns with exactly one non-zero entry and ∼75% of the rows with no non-zero entries.

#### 2.1.2 Biomedical Collaborative Graph

It is difficult to capture the complex disease and gene features in the sparse disease-gene graph, which motivates us to integrate multiple biologically networks to enrich disease-gene information. Specifically, we constructed disease-disease and gene-gene collaborative graphs. The related associations are collected from four biologically networks, including:

1. **PharmKG[15]** compose of more than 500 000 individual interconnections between genes, drugs and diseases, with 29 relation types over a vocabulary of 8000 disambiguated entities. We extracted 2406 disease-disease and 23008 gene-gene associations from PharmKG.
2. **GNBR[16]** is constructed by extracting disease, gene and chemical entities from Medline abstracts and finding connecting dependency paths between pairs of entities in single sentences. In this work, we consider the gene-gene relationship in GNBR, which contain 9 specific themes and 83335 gene-gene associations.
3. **MimMiner[6]**, a disease similarity network, is built on the combination of annotation-based methods and sequence-based methods using disease information. MimMiner has been previously used for prioritizing and contributes 18468 disease-disease associations to our collaborative graph.
4. **TTD[19]** contains message about the target disease condition, the pathway information and the corresponding drugs/ligands directed at each of these targets. In this work, we extract 3427 disease-disease and 9401 gene-gene associations from TTD.

To integrate these databases, we unified the gene names with Entrez Gene ID and the disease names according to the Medical Subject Headings (MeSH). Finally, we mapped 24301 disease-disease relationships and 115744 gene-gene relationships obtained from biologically heterogeneous networks to the disease-gene graph. Details of the statistics of these datasets are given in table 1.

**Table 1.** statistic of the dataset.

|  |  | Disease | Gene |
| --- | --- | --- | --- |
| Statistic of<br>the dataset | Number of nodes | 3209 | 12305 |
|  | Associations with same type<br>(Disease-disease, gene-gene) | 24301 | 115744 |
|  | Associations with different types<br>(Disease-gene) | 3209 | 3209 |
| Analysis of<br>the dataset | Degree with same type | 15.14 | 18.94 |
|  | Degree with different type | 1.23 | 0.32 |

### 2.2 The architecture of MiGCN

Figure 1 shows the overall framework of MiGCN. First, for each node in the disease-disease and gene-gene collaborative graphs, we initial its embedding and estimate its mutual information with the surrounding neighborhoods. Then, neighbors carrying high mutual information are selected to construct a new subset of neighbors. Node embedding and the new neighbor subset are input to the message passing function to obtain nodes final representation. Finally, we infer associations between diseases and genes from their representations. The keys of MiGCN are the mutual information estimation module and the message passing module. Details of these two modules are explained in the following sections.

**Figure 1.**
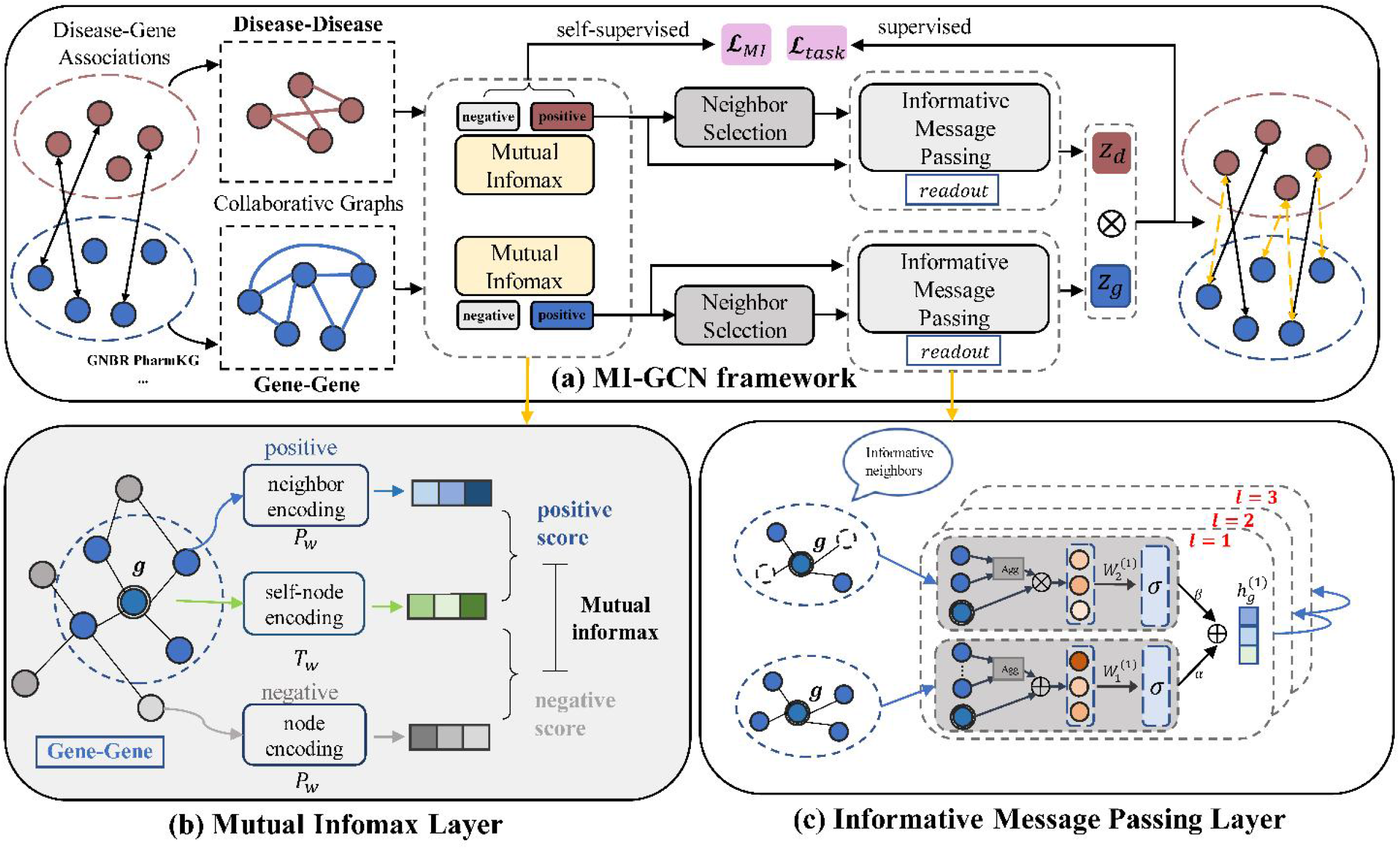
(a): Overview of MiGCN for disease-gene associations prediction. (b), (c): two key components Mutual Infomax Layer and Informative Message Passing Layer of MiGCN. The disease-disease and gene-gene collaborative graphs are input to the mutual infomax layer that estimate the mutual information between nodes and construct neighbor subset for each node. Then each node and its neighbor subset are fed into the informative message passing function to obtain the final representation.

#### 2.2.1 Graphical Mutual Infomax Layer

Given the disease-gene collaborative graph denoted as *G* = (*V* = *V*_d_ ⋃ *V*_g_, *E*), where *V*_d_, *V*_g_ represent set of diseases and genes, respectively, it is essential to parameterize nodes with graph-structured information preserved and redundant information removed. To this end, we introduce a novel graphical mutual infomax layer, constructing neighbor subset that carry high mutual information with their surrounding neighborhoods by mutual information estimation and maximization. We first calculate mutual information between with node and neighbors with negative sample strategy and then construct neighbor subset.

##### Mutual information estimation

Mutual information (MI) of two random variables is a measure of the mutual dependence between them. Mathematically, we let a random variable v to represent the node feature of node *v* ∈ *V*, then we define the probability distribution of v as:

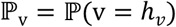

where *h*_*v*_ is the initial embedding of node *v*. Similarly, the feature distribution of the neighbor node of *v* is 𝕡_u_. We then consider the mutual information between node *v* feature and its one neighbor *u* feature which is defined as the KL-divergence between the joint distribution 𝕡_v,u_ = 𝕡(v = *h*_*v*_, u = *h*_*u*_) and the product of the marginal distributions of two random variable[20]; that is:

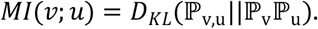

Through Donsker-Varadhan Theory[21], we can estimate the lower bound of mutual information as follow:

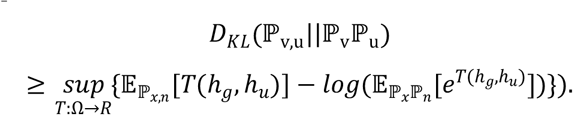

where *T* is an arbitrary function that maps the node distribution to a real number. Since it is impossible to go through all the function to evaluate the exact value of *MI*(*v*;*u*), we parameterize *T* as a neural network discriminator *D*_*w*_ (*h*_*v*_, *h*_*u*_) = *S*_*w*_(*T*_*w*_(*h*_*v*_), *P*_*w*_(*h*_*u*_)), where the subscript *w* indicates the associated functions are trainable. We implement *S*_*w*_, *T*_*w*_, *P*_*w*_ as follows:

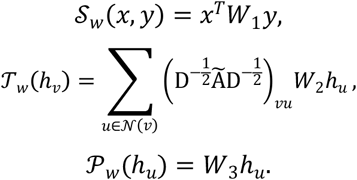

*S*_*w*_ is a score function that quantifies the affinity between node and neighbor, T_w_ is a GCN network where 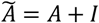 and D is the degree matrix of 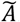. P_w_ is a fully connected network. Here, we finally define the following parameterized mutual information of node *v* and its neighbors, that is:

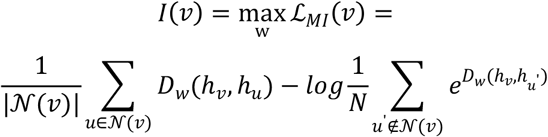

where *N* is the number of non-neighboring nodes. Since it is very consuming to include all the non-neighboring nodes, we perform negative sampling to approximate it. By training w, we can maximize the mutual information between node *υ* and its all neighbors and obtain more reasonable representation of *h*_v_.

##### Neighbor infomax selection

To eliminate noise from prior knowledge (disease and gene related interactions), we select neighbors for each node that carry high mutual information, called informative neighbors. Given a node υ and neighbor subset size *K*, we obtain the informative neighbors subset by solving:

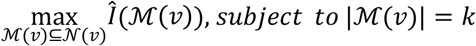

Where Î (ℳ (υ)) is a neighborhood subset criterion function that measure the ability of neighbor subset to express all neighborhoods. To solve it, we first calculate the affinity of node itself and each neighbor *u*. Here we consider the first term in mutual information:

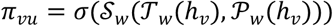

where the activation function σ we employed is sigmoid function that convert the score to the probability of being informative neighbors. Then Î (ℳ (υ)) is defined as the sum of affinity score of the neighbor subset:

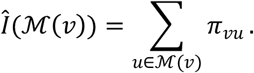

To optimize it, in practice, we calculate the affinity of node υ and all its neighbors, and sort to select the top-*K* neighbors, where *K* can be replaced as a ratio of the number of neighbors.

#### 2.2.2 Informative Message Passing Layer

The traditional graph convolution network aggregates all neighbors of a vertex and updates the vertex through a message passing function *f*:

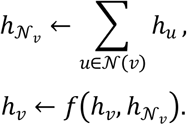

Since we obtain the informative neighbors through the mutual information calculation, namely ℳ(υ), multi-scale neighbor information can be considered as follows:

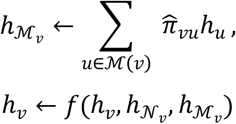

where 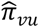 is normalized across all neighbors connected with υ by adopting the softmax function:

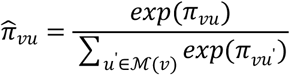

By doing so, we assign a weight to each neighbor in ℳ(υ). The next step is to design the updating function *f* which determines how to leverage the information in nodes and neighbors that is transformed to be used at next iteration. We consider the following three type variant message passing function:

##### Sub-Neighbor

Similar to the naive graph convolution network, we first consider the summation of ego representations and informative neighbor representations as follow:

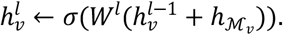

##### Attentive

The informative neighbors can be considered more important in message passing; hence it is reasonable to add more weights to them as an attention mechanism, which can be implemented by summing three representations up in different weights.

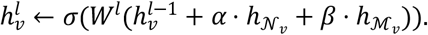

##### Informative

Another simple but useful way is that considering two kinds of feature interactions for three features which allow multi-scale neighbor information to communicate. It can be formulated as below:

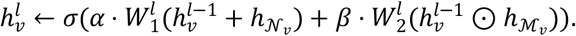

where W^*l*^ is the trainable parameter in layer *l*, α and β are hyper-parameters that control the balance of different neighbors set. After performing layer, we adopt the readout function[22] that concatenates the embeddings of each layer into a single vector:

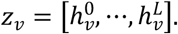

#### 2.2.3 Model Prediction and Optimization

At the end of the model, we obtain the representation of each node, specifically, the disease representation *Z*_*d*_ and the gene representation *Z*_*g*_, then the disease-gene matching score is obtained by inner product of the representation. The higher the matching score, the more likely there is a potential link between the disease and genes:

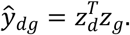

We optimize our model from two perspectives, link prediction loss and mutual information maximization loss. The former uses BPRLoss[23], which makes the matching score of known disease gene relationship pairs higher than that of unobserved disease gene pairs and it shows as follows:

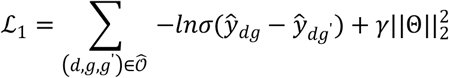

where 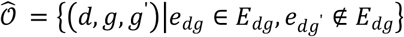, γ is the regularization coefficient, 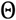 denotes all the parameters. In practice, we sample one negative pair for each observed disease-gene pair. The mutual information maximization loss is defined as ℒ_2_ = ℒ_*g*_ + ℒ_*d*_, where ℒ_*g*_ and ℒ_*d*_ are calculated as follows:

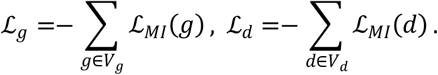

Combining the supervised and self-supervised loss, the overall loss is defined as:

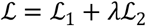

where the hyper-parameter λ balances the link prediction task and mutual information maximization task.

### 2.3 Experimental Settings

#### 2.3.1 Baselines

We compared MiGCN with three matrix-factorization (MF) based methods (BPRMF, NeuMF, IMC), two positive-unlabeled (PU) based methods (Catapult, Katz) and one GNN-based method (GCN-MF). The details of these methods are as follows:

- **BPRMF** is designed for item recommendation based on Bayesian theory to maximize the posterior probability under prior knowledge.
- **NeuMF** is a state-of-the-art factorization model, which subsumes FM under neural network.
- **IMC** combines multiple types of evidence (features) for diseases and genes to learn latent factors under the matrix factorization framework that explain the observed gene-disease associations.
- **Catapult** combines data across species using Positive-Unlabeled Learning Techniques.
- **Katz** utilizes the truncated Katz metric to construct information propagation to predict disease-gene associations.
- **GCN-MF** is the state-of-the-art approach, which employ graph convolution network to encode graph structured data. Besides, GCN-MF constructs the similarity network of disease and gene as collaborative graphs to predict disease-gene associations.

#### 2.3.2 Implementation details

We performed 5-fold cross-validation on the disease-gene training data. Each cross-validation was repeated five times, and their performance on the validation set was used to optimize all hyperparameters through grid search. The final average performance of 5-fold was reported. Specifically, we employed a two-layer message passing network with hidden units of 512 and the following set of hyperparameters: γ =0.01, λ =0.1, learning rate of 0.001, dropout rate of 0.1 and the ratio to select informative neighbors is 0.5. We implemented the proposed method with Pytorch 1.13.0 on an Nvidia GeForce RTX 3090 GPU.

#### 2.3.3 Evaluation

Similar to previous study, the accuracy of the model is measured by three widely used evaluation protocols: recall@N, precision@N and AUC:

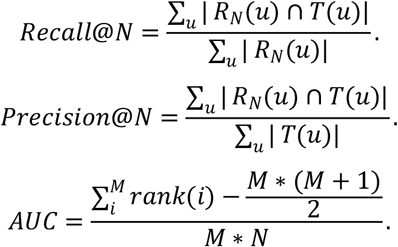

where *R*_*N*_(*u*) denotes the set of nodes predicted to be associated with *u, T*(*u*) denotes the set of nodes associated with *u* in the test set, M, N represent the number of positive and negative samples, *rank*(*i*) denotes the position of the i-th positive sample score after ranking.

For each disease in the test set, we calculate its score ŷ with all genes, except the genes associated with the disease in the training set. The genes with the top-N highest score were selected as the predictive associated genes set *(R*_*N*_(*u*)). In our work, we select N in {1,5,10,15}. For each metric, we first compute the results of each disease in the test set and report the average metric

## 3 Results and discussion

### 3.1 MiGCN outperforms other competing methods

We evaluated the proposed MiGCN on OMIM datasets and compared the performance with other state-of-the-art methods. Following the prior work[11], we conduct 5-fold cross-validation (CV) and report their metric scores’ mean (Table 2) and standard deviation (Table S1 and S2). As shown in table 2, MiGCN outperforming all existing methods including MF-based and graph-based method. Specifically, MiGCN achieves AUROC values of 0.831 which improves over the strongest baseline by 8.63%. As the dataset is extremely imbalanced, we also report the precision and recall of top-n predictions, where MiGCN achieves recall@15 and precision@15 of 0.2194 and 0.0169, with 55.27% and 55.14% improvement over the strongest baselines, respectively. The consistent improvement demonstrated the effectiveness of MiGCN.

**Table 2.**
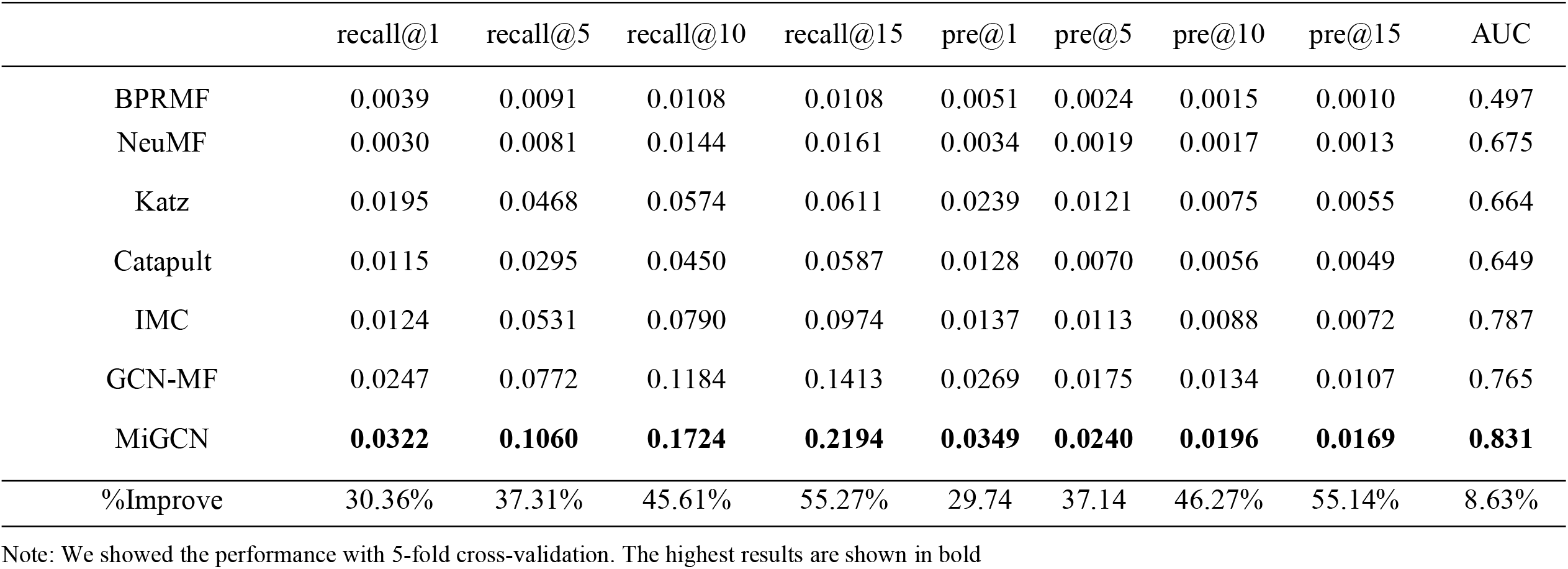
Performance comparison of MiGCN with state-of-the-art methods.

|  | recall@1 | recall@5 | recall@10 | recall@15 | pre@1 | pre@5 | pre@10 | pre@15 | AUC |
| --- | --- | --- | --- | --- | --- | --- | --- | --- | --- |
| BPRMF | 0.0039 | 0.0091 | 0.0108 | 0.0108 | 0.0051 | 0.0024 | 0.0015 | 0.0010 | 0.497 |
| NeuMF | 0.0030 | 0.0081 | 0.0144 | 0.0161 | 0.0034 | 0.0019 | 0.0017 | 0.0013 | 0.675 |
| Katz | 0.0195 | 0.0468 | 0.0574 | 0.0611 | 0.0239 | 0.0121 | 0.0075 | 0.0055 | 0.664 |
| Catapult | 0.0115 | 0.0295 | 0.0450 | 0.0587 | 0.0128 | 0.0070 | 0.0056 | 0.0049 | 0.649 |
| IMC | 0.0124 | 0.0531 | 0.0790 | 0.0974 | 0.0137 | 0.0113 | 0.0088 | 0.0072 | 0.787 |
| GCN-MF | 0.0247 | 0.0772 | 0.1184 | 0.1413 | 0.0269 | 0.0175 | 0.0134 | 0.0107 | 0.765 |
| MiGCN | **0.0322** | **0.1060** | **0.1724** | **0.2194** | **0.0349** | **0.0240** | **0.0196** | **0.0169** | **0.831** |
| %Improve | 30.36% | 37.31% | 45.61% | 55.27% | 29.74 | 37.14 | 46.27% | 55.14% | 8.63% |

Note: We showed the performance with 5-fold cross-validation. The highest results are shown in bold

Among the baselines, the naive MF-based methods (BPRMF and NeuMF) achieve the poorest performance, which may be ascribed to the poor ability of low rank matrix approximation to capture the nonlinear features in graph data. By contrast, IMC performs better by combining the disease description and gene expression features. However, the data-driven approach IMC suffers from the high cost of generating reasonable features, while our model outperforms it with random initial features. Compared with IMC, GCN-MF achieves higher recall and precision but lower AUROC. One possible reason is that GCN-MF only learns local information of first-order neighbors so that it cannot well predict the long-distance node relationship. However, increasing the hop number of neighbors would bring redundant and harmful information. Notably, MiGCN is able to consider high-order neighbors and eliminate noise through mutual information estimation and informative neighbor selection.

### 3.2 MiGCN is able to effectively eliminate noise

The key contribution of our model is the ability to eliminate noise in graphs. Firstly, to investigate the influence of the noise introduced by prior knowledge, we replaced our collaboration graphs with similarity graphs constructed by GCN-MF and re-trained our model, termed MiGCN_sim_. The similarity graphs include disease-disease and gene-gene similarity, which are hand-crafted and contain more unpredictable noise. As shown in figure 2, the performance of GCN-MF decreases dramatically as GCN layers stacking, due to their poor ability to eliminate noise from high-order neighbors. By contrast, our model is able maintain stable and consistently outperforms GCN-MF on AUC with a large margin. With further stacking, the performance of the model starts to descend gradually which is expected as a result of over-fitting.

**Figure 2.**
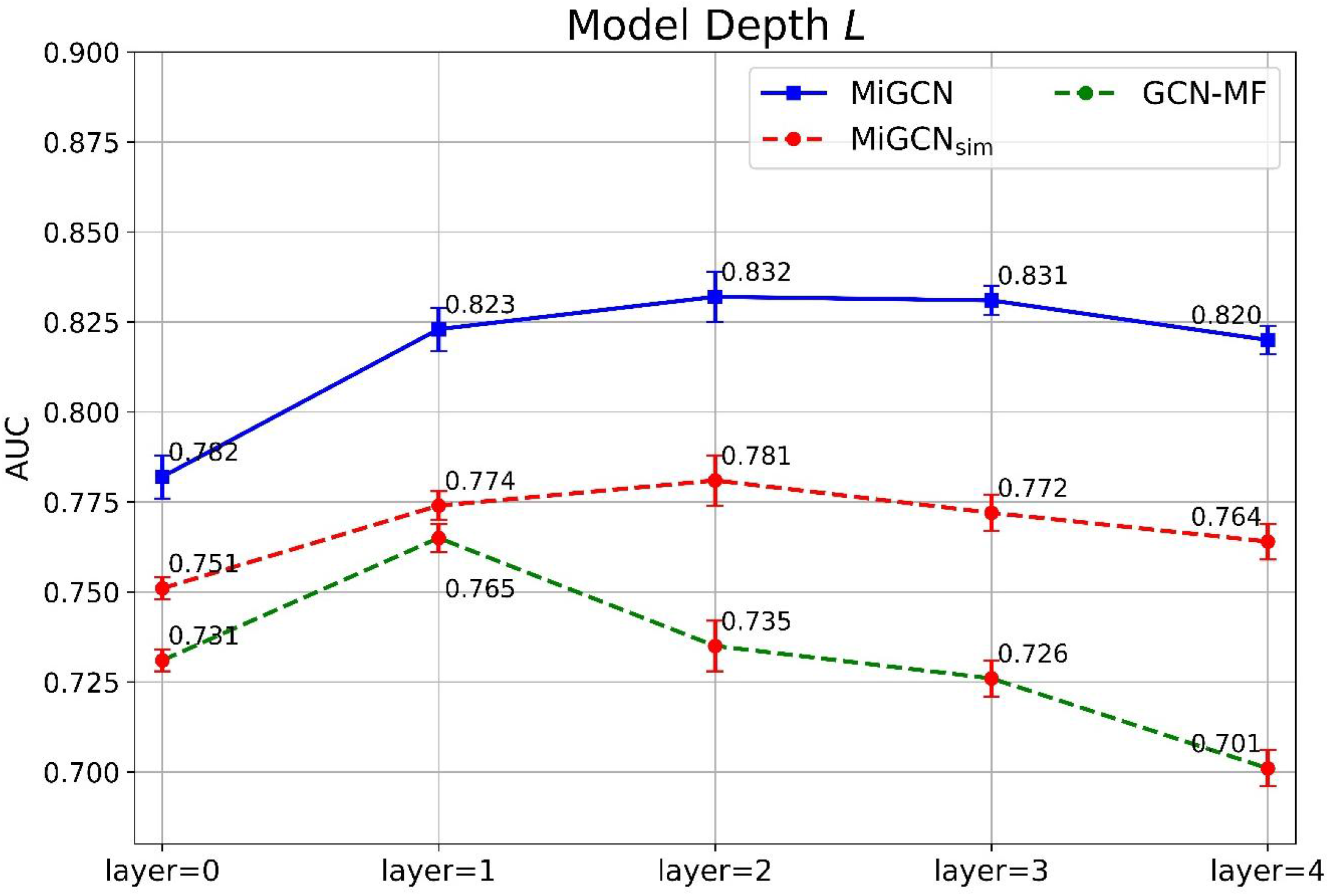
Comparison between MiGCN and GCN-MF on similarity graphs with different model depth

In addition, message passing function is a significant factor for noise elimination which could directly influence information flow. To explore how the message passing function affects the performance, we compared our proposed informative message passing function with other variants (sub-neighbor-based, attentive-based, and GCN-based). As shown in figure 3, the informative message passing function achieves the best performance both on MiGCN and MiGCN_sim_. When replacing the informative function on MiGCN with GCN and sub-neighbor-based function, the model showed relatively low performance with recall@15 of MiGCN dropping by 1.6% and 2.4%, respectively. Particularly, the recall@15 decreases by more than 9% on MiGCN_sim._ These results indicate that the performance of the model is affected by the noise and demonstrate the rationality and effectiveness of informative message passing. The attentive-based function is slightly inferior to the informative function but superior to sub-neighbor and GCN-based function. It shows that the attention mechanism also eliminates the noisy information from neighbors, but the informative function further incorporates additional feature interaction and thus improve information flowing.

**Figure 3.**
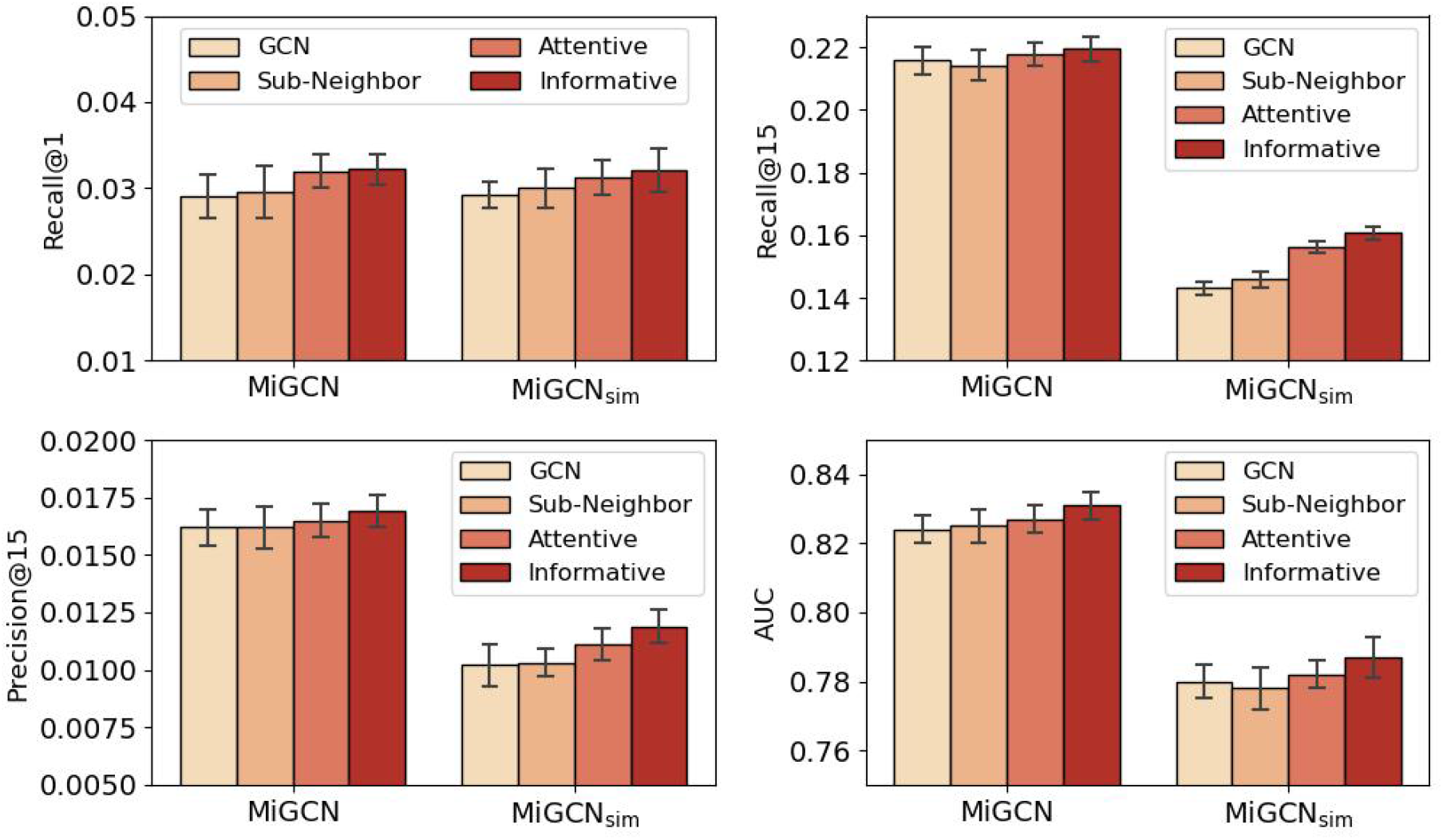
Effect of Message Passing Function.

### 3.2 Model components analysis

In order to study the contributions of each module of MiGCN, we provided ablation studies in this section. As shown in table 3, the removal of the mutual infomax module (w/o MI) caused a decrease of 2%, 3%, and 1% in terms of recall@15, precision@15 and AUC, respectively. This change indicated that the mutual infomax module can effectively remove noises from the data and thus improve the performance. The removal of the informative message passing module (w/o MP) also causes degradation to the model performance, which indicated that the information neighbor subset contributes to a richer flow of information between nodes, thus improving the learning ability of the GCN. When the two modules are both excluded, the model performed the worst, with recall@15, precision@15 and AUC dropping by 3%, 6% and 1.6%, respectively. In summary, the better performance of our model relied on the cooperation of these modules.

**Table 3.**
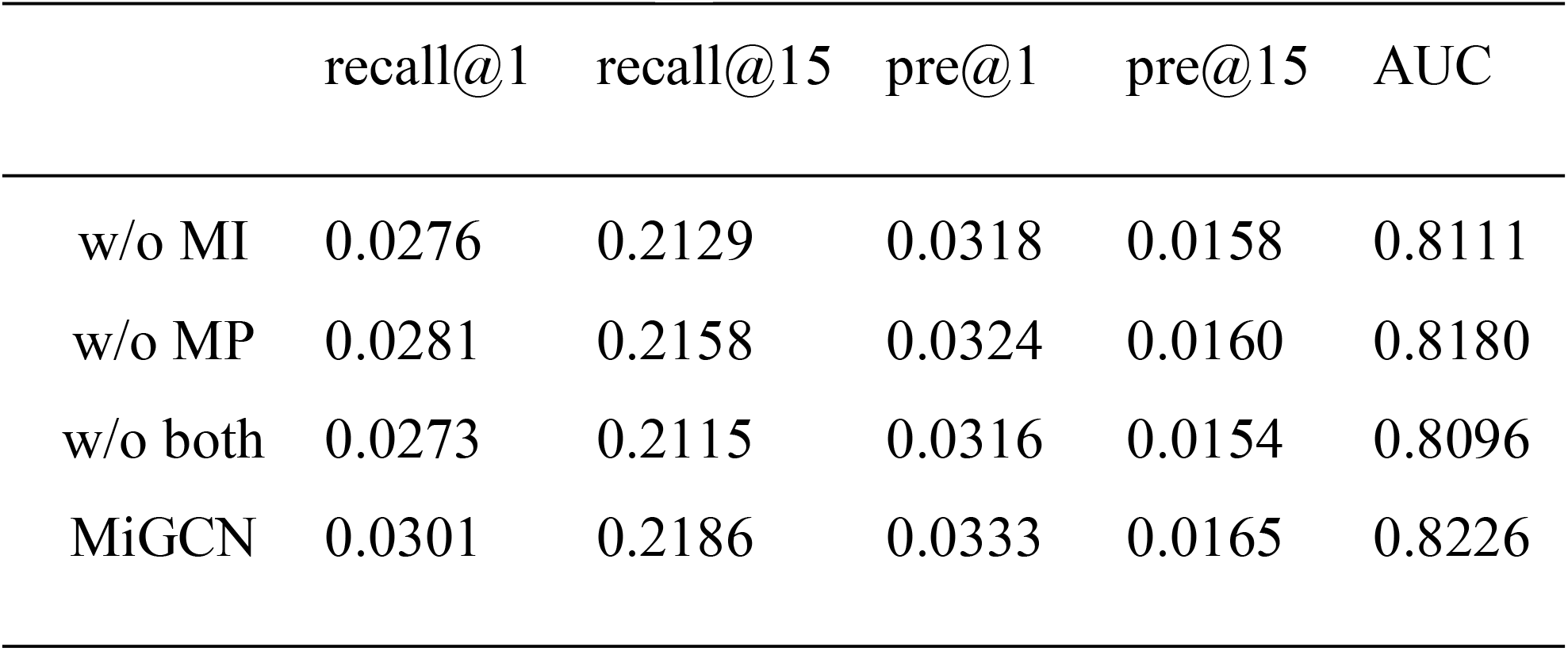
Ablation study on different module of MiGCN.

### 3.3 Visualization of MiGCN

To gain insights into how the final embeddings represent the gene and the disease features, we obtained the nodes representation learned by three model: GCN-MF, MiGCN_sim_ and MiGCN, and mapped into a 2-dimensional space using t-SNE[24] for visualization. Specifically, we extracted representations of disease-gene pairs by element-wise product of their respective final embeddings, and then projected all pairs into 2D space. The associated disease-gene pairs are colored in blue while the non-associated pairs are colored in pink. As shown in figure 4, there is a clear boundary between the associated and unassociated pairs learned from MiGCN, while GCN-MF cannot well separate them. MiGCN_sim_ can also separate the associated and non-associated disease-gene pairs, but there are larger overlapping parts compared with MiGCN.

**Figure 4.**
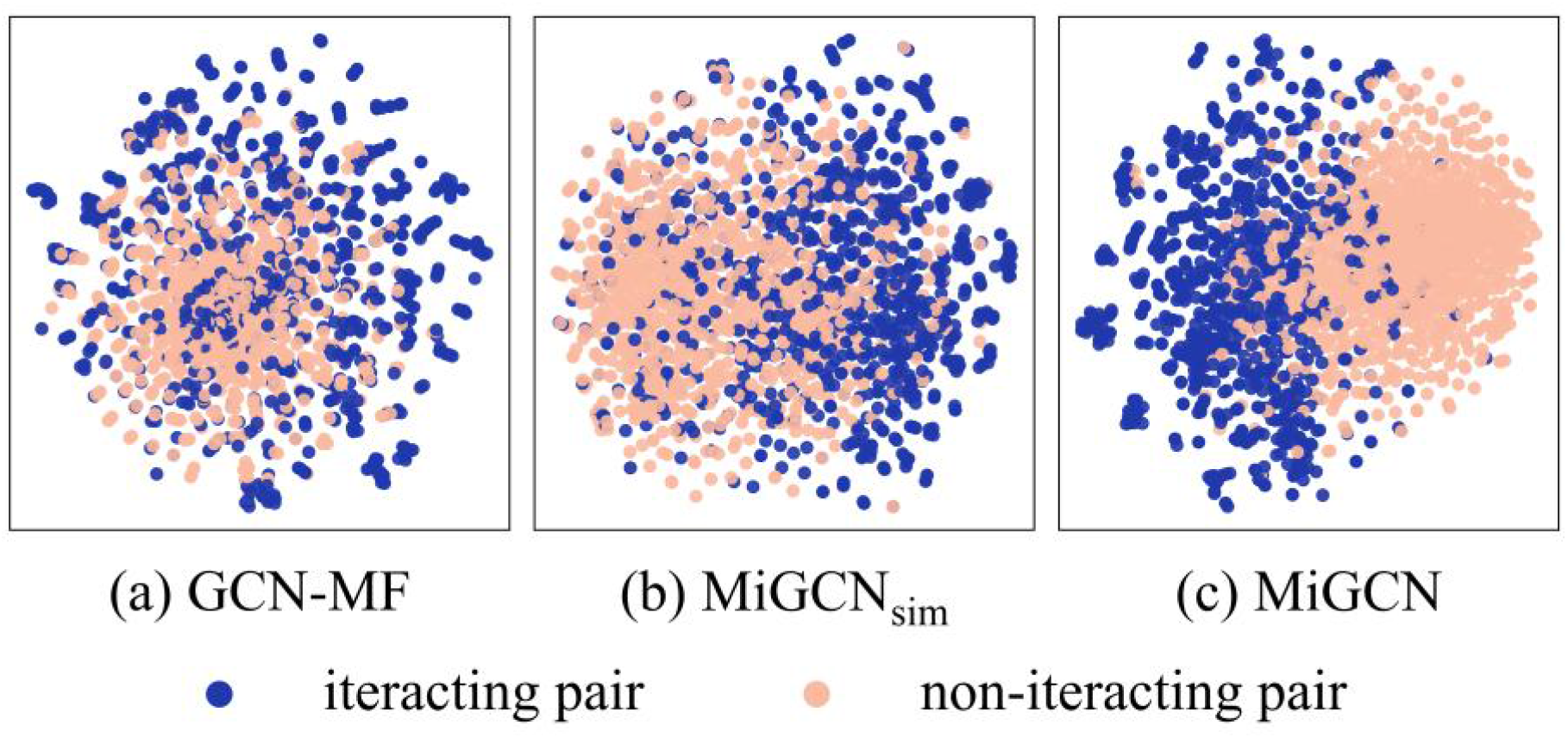
Projection of associated (blue) and unassociated (pink) disease-gene pairs

### 3.4 Case study of MiGCN

To further validate the prediction ability of MiGCN, we conducted a case study to predict associated genes with one of the most common diseases: *Alzheimer*. Table 4 shows the novel disease-gene associations, with disease, gene name and literature reference supporting interpretation. For *Alzheimer (AD)*, among the top 10 predicted disease-gene associations ranked according to their prediction scores, four genes were not included in our dataset but indicated associated from recenter literature. For instance, the prion protein gene (PRNP) is predicted to associated with AD. A previous study[25] reported that the association between V homozygosity at PRNP codon 129 and cognitive performance in the elderly. The meta-analysis[26] also revealed that subjects homozygous at codon 129 had a 1.3-fold increased risk of developing AD compared to heterozygous individuals. Figure 5 shows the deduction for the predicted *Alzheimer -PRNP* association. The path was inferred by a 3-hopper through their closest neighbors (*Pancreatitis* and *SPINK1*) that were measured by the estimated mutual information scores, which may provide evidence for the underlying mechanisms of the disease-gene association.

**Table 4.** Top-10 associations of Alzheimer predicted by MiGCN.

| Top-N | Associated genes | Predicting scores | Supporting evidence |
| --- | --- | --- | --- |
| 1 | PRNP | 3.60 | [25, 26] |
| 2 | TRPM7 | 3.44 |  |
| 3 | GRN | 3.43 | [27] |
| 4 | PSEN1 | 3.24 | [28] |
| 5 | CST3 | 3.14 |  |
| 6 | HLA-DQB1 | 3.14 |  |
| 7 | APOA1 | 3.14 |  |
| 8 | SORL1 | 3.12 | OMIM[3] |
| 9 | APOE | 3.11 | [29] |
| 10 | SERPINI1 | 2.99 |  |

**Figure 5.**
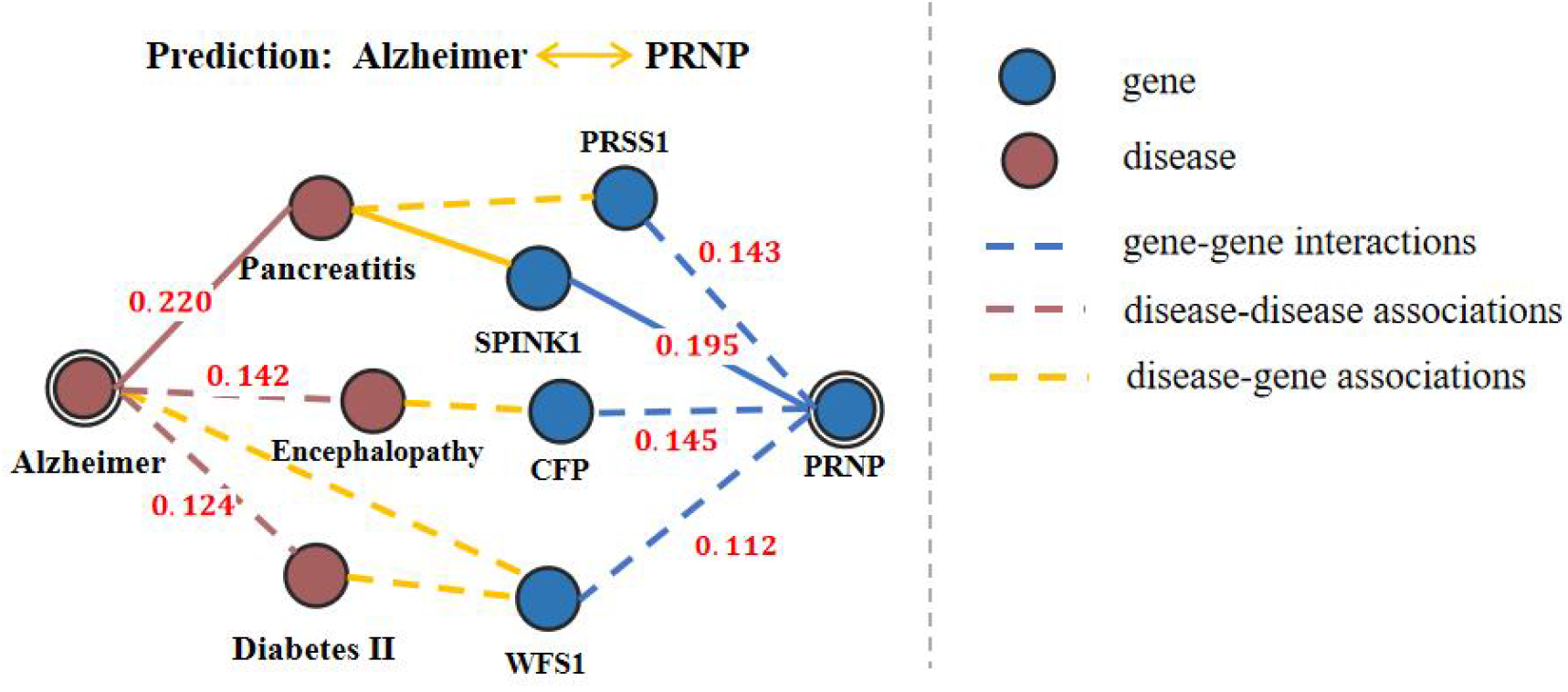
Real example of novel disease-gene association predicted by MiGCN.

Two other common disease *Parkinson* and *Diabetes* are also selected to explore the predictive performance of the MiGCN. The top 10 predicted genes are list in supplementary table S3 and S4. For *Parkinson*, seven of ten (70%) predicted genes were validated by previous studies from literature. For *Diabetes*, five of ten (50%) predicted genes are found supported by evidence from the literature. These results again indicate the generalization ability of our model.

## 4 Discussion

Discovering disease-gene association is a fundamental and critical biomedical task, which assists biologists and physicians to discover pathogenic mechanism of syndromes. Existing MF-based and graph-based methods predict disease-gene associations by combining rich prior knowledge of diseases and genes, but both suffer from the inevitable noise from the prior knowledge. In this work, we propose a new framework MiGCN to address the problem of disease-gene associations prediction. We first construct two high quality disease-disease and gene-gene collaborative graphs by integrating multiple biologically networks. To explore and exploit them, MiGCN incorporates self-supervised and supervised learning, where the self-supervised learning selects high-quality neighbors and optimizes node representations by maximizing mutual information between nodes and neighbors, and the supervised learning updates node representations and predicts disease-gene associations through a newly designed informative message passing combining informative neighbors subset. Experiments on the public dataset demonstrate the high performance of MiGCN and highlight its ability to eliminate noise from the external knowledge source.

However, there is still room for further improvements on MiGCN. First of all, although we made efforts to assemble and utilize external biological networks, we did not consider the heterogeneity of these networks i.e., the node and edge types are ignored in our framework. The predictive performance could be improved if node and edge types information can be used effectively. Secondly, knowledge graph techniques could be applied on the collaborative graphs to optimize entity representation.

In general, this work explores the potential of graph neural networks in association prediction. We find that combining diverse biomedical sources and learning models could obtain large improvement of predictive performance. This method can also be used to infer other associations, such as gene-drug, disease-drug, and drug-drug relationships. In the future, we would integrate more biomedical knowledge sources to further improve our model.

## Key points

- We constructed two high quality disease-disease and gene-gene collaborative graphs by combing four biological networks to enrich the sparse disease-gene information, and proposed a new GCN-based model, MiGCN, by using a self-supervised mutual infomax module to eliminate noise and a supervised message passing module to predict disease-gene associations.
- We introduced an enhanced informative message passing mechanism to strengthen the role of informative neighbors during GCN learning.
- We conducted extensive experiments to demonstrate the effectiveness of MiGCN that outperforms the state-of-the-art method with a large margin.

## Data availability

The datasets, source codes and trained models of MiGCN are available at https://github.com/biomed-AI/MiGCN.

## Supplementary data

Supplementary data are available online at https://academic.oup.com/bib.

*Conflict of Interest:* none declared.

## References

1. Denny, J.C., et al., PheWAS: demonstrating the feasibility of a phenome-wide scan to discover gene–disease associations. 2010. 26(9): p. 1205–1210.

2. Özgür, A., et al., Identifying gene-disease associations using centrality on a literature mined gene-interaction network. 2008. 24(13): p. i277–i285.

3. Hamosh, A., et al., Online Mendelian Inheritance in Man (OMIM), a knowledgebase of human genes and genetic disorders. 2005. 33(suppl_1): p. D514–D517.

4. Yang, P., et al., Positive-unlabeled learning for disease gene identification. 2012. 28(20): p. 2640–2647.

5. Mordelet, F. and J.-P.J.B.b. Vert, Prodige: Prioritization of disease genes with multitask machine learning from positive and unlabeled examples. 2011. 12(1): p. 1–15.

6. Adie, E.A., et al., SUSPECTS: enabling fast and effective prioritization of positional candidates. 2006. 22(6): p. 773–774.

7. Sirota, M., et al., Discovery and preclinical validation of drug indications using compendia of public gene expression data. 2011. 3(96): p. 96ra77–96ra77.

8. Lee, I., et al., Prioritizing candidate disease genes by network-based boosting of genome-wide association data. 2011. 21(7): p. 1109–1121.

9. Natarajan, N. and I.S.J.B. Dhillon, Inductive matrix completion for predicting gene–disease associations. 2014. 30(12): p. i60–i68.

10. Singh-Blom, U.M., et al., Prediction and validation of gene-disease associations using methods inspired by social network analyses. 2013. 8(5): p. e58977.

11. Han, P., et al. GCN-MF: disease-gene association identification by graph convolutional networks and matrix factorization. in Proceedings of the 25th ACM SIGKDD international conference on knowledge discovery & data mining. 2019.

12. Kipf, T.N. and M.J.a.p.a. Welling, Semi-supervised classification with graph convolutional networks. 2016.

13. Song, Y., et al. Communicative Representation Learning on Attributed Molecular Graphs. in IJCAI. 2020.

14. Mai, S., et al. Communicative Message Passing for Inductive Relation Reasoning. in AAAI. 2021.

15. Zheng, S., et al., PharmKG: a dedicated knowledge graph benchmark for bomedical data mining. 2021. 22(4): p. bbaa344.

16. Percha, B. and R.B.J.B. Altman, A global network of biomedical relationships derived from text. 2018. 34(15): p. 2614–2624.

17. Velickovic, P., et al., Deep Graph Infomax. 2019. 2(3): p. 4.

18. Peng, Z., et al. Graph representation learning via graphical mutual information maximization. in Proceedings of The Web Conference 2020. 2020.

19. Chen, X., Z.L. Ji, and Y.Z.J.N.a.r. Chen, TTD: therapeutic target database. 2002. 30(1): p. 412–415.

20. Belghazi, M.I., et al. Mutual information neural estimation. in International conference on machine learning. 2018. PMLR.

21. Donsker, M.D., S.S.J.C.o.P. Varadhan, and A. Mathematics, Asymptotic evaluation of certain Markov process expectations for large time, I.m 1975. 28(1): p. 1–47.

22. Xu, K., et al. Representation learning on graphs with jumping knowledge networks. in International conference on machine learning. 2018. PMLR.

23. Zhang, F., et al. Collaborative knowledge base embedding for recommender systems. in Proceedings of the 22nd ACM SIGKDD international conference on knowledge discovery and data mining. 2016.

24. Van der Maaten, L. and G.J.J.o.m.l.r. Hinton, Visualizing data using t-SNE. 2008. 9(11).

25. Del Bo, R., et al., Is M129V of PRNP gene associated with Alzheimer’s disease? A case-control study and a meta-analysis. 2006. 27(5): p. 770. e1–770. e5.

26. Berr, C., et al., Polymorphism of the prion protein is associated with cognitive impairment in the elderly: the EVA study. 1998. 51(3): p. 734–737.

27. Jin, S.C., et al., Pooled-DNA sequencing identifies novel causative variants in PSEN1, GRN and MAPT in a clinical early-onset and familial Alzheimer’s disease Ibero-American cohort. 2012. 4(4): p. 1–9.

28. Lanoiselée, H.-M., et al., APP, PSEN1, and PSEN2 mutations in early-onset Alzheimer disease: A genetic screening study of familial and sporadic cases. 2017. 14(3): p. e1002270.

29. Genin, E., et al., APOE and Alzheimer disease: a major gene with semi-dominant inheritance. 2011. 16(9): p. 903–907.

